# Towards the Clinical Translation of a Silver Sulfide Nanoparticle Contrast Agent: Large Scale Production with a Highly Parallelized Microfluidic Chip

**DOI:** 10.1101/2023.12.02.569706

**Authors:** Katherine J. Mossburg, Sarah J. Shepherd, Diego Barragan, Nathaniel H. O, Emily K. Berkow, Portia S. N. Maidment, Derick N. Rosario Berrios, Jessica C. Hsu, Michael J. Siedlik, Sagar Yadavali, Michael J. Mitchell, David Issadore, David P. Cormode

## Abstract

Ultrasmall silver sulfide nanoparticles (Ag_2_S-NP) have been identified as promising contrast agents for a number of modalities and in particular for dual-energy mammography. These Ag_2_S-NP have demonstrated marked advantages over clinically available agents with the ability to generate higher contrast with high biocompatibility. However, current synthesis methods are low-throughput and highly time-intensive, limiting the possibility of large animal studies or eventual clinical use of this potential imaging agent. We herein report the use of a scalable silicon microfluidic system (SSMS) for the large-scale synthesis of Ag_2_S-NP. Using SSMS chips with 1 channel, 10 parallelized channels, and 256 parallelized channels, we determined that the Ag_2_S-NP produced were of similar quality as measured by core size, concentration, UV-visible spectrometry, and *in vitro* contrast generation. Moreover, by combining parallelized chips with increasing reagent concentration, we were able to increase output by an overall factor of 3,400. We also found that *in vivo* imaging contrast generation was consistent across synthesis methods and confirmed renal clearance of the ultrasmall nanoparticles. Finally, we found best-in-class clearance of the Ag_2_S-NP occurred within 24 hours. These studies have identified a promising method for the large-scale production of Ag_2_S-NP, paving the way for eventual clinical translation.

## Introduction

Inorganic nanoparticles are being explored for many biomedical applications such as imaging and have been identified as a valuable option for improving disease detection.^1-4^ Their tunable properties, such as photon attenuation, size, shape, and surface charge, allow them to be adapted to a wide variety of imaging modalities and disease settings.^5-7^ Thus, much attention has been focused on designing inorganic nanoparticles for improved cancer detection and monitoring. Recent work has identified sub-5nm glutathione-coated silver-sulfide nanoparticles (Ag_2_S-NP) as a promising imaging contrast agent that is especially well-suited for breast cancer screening due to their strong x-ray attenuation in the energy range used for mammography.^8-10^ Ag_2_S-NP have demonstrated improved contrast over traditional iodinated agents and have a longer circulation window.^8^ Furthermore, they can be synthesized at room temperature, under ambient conditions, and in water, while other inorganic nanoparticles require high temperatures and toxic reagents for synthesis.^2, 11-15^ Additionally, silver-based nanomaterials are much less expensive than their gold counterparts, making them more practical economically.^16^ Previous studies have shown that Ag_2_S-NP also are highly biocompatible.^8-10, 17^ Due to their ultrasmall size, these Ag_2_S-NP are renally excretable, which is an improvement over many alternative inorganic nanoparticle formulations and makes them appropriate for eventual clinical translation.^7, 18^-^20^ However, to facilitate large animal studies and clinical translation of Ag_2_S-NP, it is necessary to have a rapid and reliable synthesis method.

Current synthesis methods available for Ag_2_S-NP either produce nanoparticles of variable physical characteristics (bulk synthesis) or in quantities that are too small for clinical relevance (microfluidics).^21-23^ Bulk synthesis methods are one option for upscaling to larger volumes, but suffer from temperature and concentration gradients that result in nanoparticles with heterogenous size distributions. Additionally, because bulk methods rely on batch processes, there is variability amongst various runs of the synthesis. Microfluidic mixing techniques, such as staggered herringbone mixing, are highly valuable tools for decreasing dispersity, controlling size, and ensuring reproducibility compared to bulk mixing; however, they have been limited by low throughputs.^24-27^ The previously reported microfluidic synthesis method for Ag_2_S-NP produces only about 20 mg of product (by silver mass) per hour, which then must be concentrated and purified.^10^ At this rate, it would take over two weeks to produce one dose for the average American woman for the synthesis alone, based on a moderate dose of 100 mg Ag/kg.^28^

To overcome the current limitations of large-scale nanoparticle production, this study sought to evaluate Ag_2_S-NP production using a scalable silicon microfluidic system (SSMS) to confirm that nanoparticles are of the same quality as those produced on a small scale. SSMS is a novel microfluidic platform which allows for solutions to flow through multiple parallelized staggered herringbone mixing channels simultaneously. In this study, we focused on SSMS with 1 channel, 10 parallelized channels, and 256 parallelized channels (Figure 1). With only a single set of inputs and outputs, for delivery of reagents and collection of nanoparticles, respectively, the SSMS allows for the logistical ease of operating only one device with the production power of many. The unique design of this platform includes flow resistors which ensure equal flow rates in each parallel channel, resulting in identical product synthesis across the chip.^29^ The materials used in the fabrication of SSMS were selected to meet industry standards for pharmaceutical production, paving the way for eventual translation.^30^ Previous work with a system similar to SSMS has indicated that it is possible to produce high-quality nanomaterials at a drastically increased rate, but these studies were done with organic nanoparticles.^29, 31-34^ With a microfluidic device designed for large-scale production of SSNP, Ag_2_S-NP produced will be more consistent and have high production rates to enable clinical studies necessary for translation to patient care. Therefore, the studies presented here are on the forefront of advancing microfluidic devices for clinical scale production of inorganic nanoparticles.

**Figure 1.**
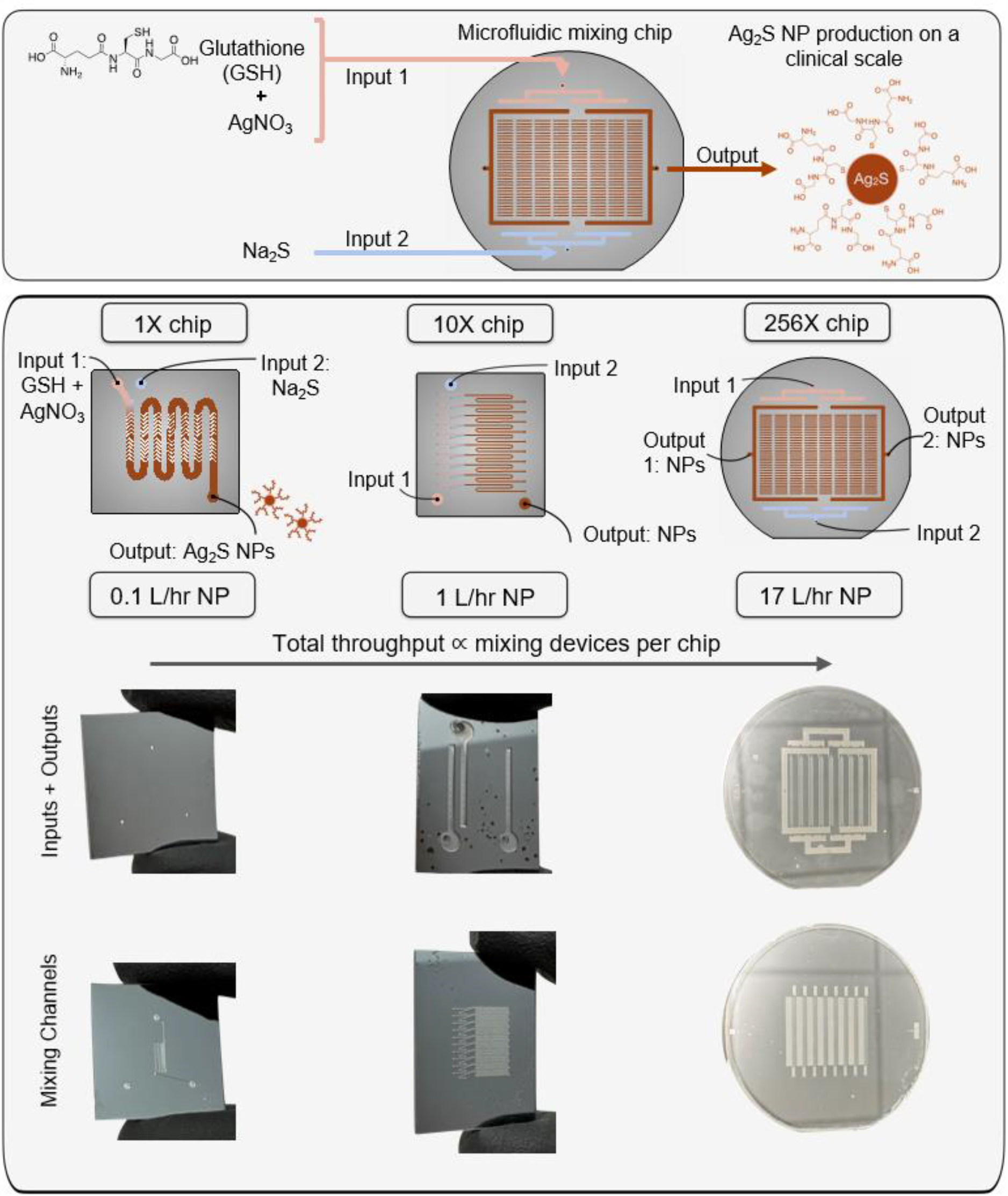
Schematic and photographs showing microfluidic chips used for synthesis of Ag_2_S-NP.

Herein, we report the rapid synthesis of monodispersed, sub-5 nm Ag_2_S-NP using the SSMS microfluidic chip with up to 256 parallel channels while monitoring the chip and the product characteristics. We evaluated the nanoparticle quality across all of the microfluidic synthesis methods through the size and absorption spectra while increasing the production rate from 0.1 L/hr to 17 L/hr. Moreover, we studied the effect of increasing reagent concentrations on Ag_2_S-NP characteristics and yield. The nanoparticles were also studied in phantom and mouse models for their contrast generation and compared to previous studies and clinically available agents. Additionally, the biodistribution of remaining silver in mouse models was measured to determine 24-hour clearance rates of Ag_2_S-NP from each synthesis method.

## Materials

Silver nitrate (AgNO_3_, 99%), sodium sulfide (Na_2_S), and L-glutathione (GSH, 98%) were purchased from Sigma-Aldrich (St. Louis, MO). Sodium hydroxide (NaOH) was purchased from Fisher Scientific (Pittsburgh, PA). A commercially available staggered herringbone single channel microfluidic chip was purchased from Microfluidic ChipShop (Jena, Germany). Iopamidol was obtained from Bracco Diagnostics (Monroe, NJ). Nude mice were acquired from The Jackson Laboratory (Bar Harbor, ME).

## Methods

### Fabrication of SSMS Chips

Microfluidic chips were fabricated in silicon and glass substrates as previously described^35^ using a process of four-layer lithography combined with ion etching to fabricate 3D structures. This 3D architecture distributes the input reagents uniformly to each individual mixing unit where the nanoparticles are formulated, then the nanoparticles are collected in a pooled output. Briefly, a single 100 mm double-sided polished silicon wafer (University Wafer, South Boston, MA) is lithographically patterned with S1805 photoresist to define the features, then etched to the intended etch depth using deep reactive ion etching. This process of lithography/etching is repeated for all four layers. The etched silicon wafer is anodically bonded to glass wafers on each side to encapsulate the microfluidic channels. Inputs and outputs are connected to the microchannels by machined holes patterned in one of the glass wafers. The bonded chip is fabricated in the Quattrone Nanofabrication Facility at the University of Pennsylvania.

To operate the silicon and glass chips, a custom pressure-driven flow system was engineered as previously described.^35^ Briefly, the two inputs are housed in pressurized vessels which are connected to a nitrogen tank and controlled by pressure controllers to regulate the input flow rates. The silicon and glass chip is encased in an aluminum housing system to connect to the input/output tubings. Chip performance is monitored by a DM4 B upright microscope (Leica Microsystems GmbH, Germany) to ensure a 3:1 flow rate ratio between the GSH + AgNO_3_ and Na_2_S inputs, respectively.

### Synthesis of Ag_2_S-NP

Ag_2_S-NP were synthesized according to a protocol adapted from previous reports.^8, 10^ The basic protocol is as follows: two solutions were prepared and co-injected into a staggered herringbone microfluidic mixing chip, either commercially purchased or fabricated as described previously. One solution was prepared with 767 mg GSH and 42.5 mg AgNO_3_ in 75 mL of deionized water, then the pH of this solution was adjusted to 7.4 using NaOH. For the second solution, 10 mg of Na_2_S was dissolved in 25 mL of water. The two solutions were co-injected into a mixing chip at a 3:1 rate, respectively, with an overall flow rate of 2 mL/min/channel. Due to limitations of the experimental setup, the SSMS 256X chip was operated at a total flow rate of 17 L/hr (=1.1 mL/min/channel). The output was collected and then concentrated and washed with DI water using 3 kDa molecular weight cut-off (MWCO) filtration tubes (Sartorius Stedim Biotech, Germany). The tubes were centrifuged at 4000 rpm in 20-minute cycles until the final output was concentrated 100-fold The concentrated nanoparticles were then filtered through a 0.020 μm filter membrane (Whatman Anotop, Boston, MA) and stored at 4°C until use in experiments.

The CS-Ag_2_S-NP were prepared according to the protocol described here using the Fluidic 187 Micro Mixer from the Microfluidic ChipShop (Jena, Germany). 1X-Ag_2_S-NP, 10X-Ag_2_S-NP, and 256X-Ag_2_S-NP were each prepared using the 1X, 10X, and 256X SSMS chips respectively. Bulk-Ag_2_S-NP were synthesized by combining the two previously described solutions in a beaker and allowing the mixture to stir for 20 minutes. After this, the solution was concentrated, washed, and filtered as described above.

Increased reagent concentrations were also used for the synthesis of Ag_2_S-NP. For these studies, the same volume of water was used to prepare each solution, however the reagent concentrations were increased to 100%, 500%, 1000%, and 2000% of the originally stated concentrations. The synthesis was then run and followed by concentration and purification as previously described.

### UV/Visible Absorption

All Ag_2_S-NP were characterized using a Thermo Scientific Genesys UV/Visible Spectrophotometer to record the absorbance of each sample. Samples were prepared by diluting Ag_2_S-NP stock solutions in deionized water to 10 μg/mL and adding 1 mL to a cuvette for analysis.

### Transmission Electron Microscopy

Each sample was analyzed using a T12 cryo-electron microscope (FEI Tecnai) operated at 100 kV. Samples were prepared by dropping 10 μL of diluted samples of Ag_2_S-NP onto formvar-coated, carbon-film stabilized copper grids (Electron Microscopy Sciences, Hatfield, PA) and allowing them to air dry. ImageJ (National Institutes of Health, Bethesda, MD) was used to measure the core diameter of the nanoparticles. 100 nanoparticles per sample were analyzed and measurements of core diameter are presented as mean ± SEM. A one-way ANOVA test with Turkey’s multiple comparisons was used to determine if nanoparticle core size was significantly different among synthetic methods.

### Inductively Coupled Plasma Optical Emission Spectroscopy

The concentration of silver in each sample was measured using Spectro-Genesis inductively coupled plasma – optical emission spectroscopy (ICP-OES). Samples were prepared by dissolving 10 μL of Ag_2_S-NP in 1 mL of nitric acid and then diluting with deionized water to a final volume of 10 mL. Each Ag_2_S-NP solution had samples prepared in triplicate.

### Phantom Imaging

Samples for phantom imaging of each type of contrast agent were prepared in triplicate with concentrations of 0, 0.5, 1, 2, 4, 6, 8, and 10 mg/mL. Each sample was loaded into a 0.2 mL tubes, placed in a rack, and covered in parafilm. Each rack of samples was scanned using a MILabs micro-CT scanner with tube voltage of 55 kV and isotropic 100 micron voxels. [matrix size = 512 x 512, field of view = 37 x 37 cm, reconstructed slice thickness = 0.5 mm]. Attenuation rates for each contrast agent were determined using Osirix MD software and a one-way ANOVA with Turkey’s multiple comparisons test was used to determine statistical significance.

### Animal Model

*In vivo* imaging was performed using 8 week-old female nude mice (Jackson Laboratories, Bar Harbor, ME) with 5 mice per group in accordance with protocols approved by the Institutional Animal Care and Use Committee of the University of Pennsylvania. Mice were scanned preinjection, then administered Ag_2_S-NP at a dose of 250 mg Ag/kg body weight of the mouse via the tail vein. Mice were anesthetized for scans using isoflurane.

### *In Vivo* Imaging

All mice were imaged using a MILabs micro-CT scanner (tube voltage = 55 kV) with scans performed at 5, 30, 60, 120, and 1440 minutes post-injection. Parameters for the scans were as follows: matrix size = 512 x 512, field of view = 37 x 37 cm, reconstructed slice thickness = 0.5 mm. Scans were analyzed using Osirix MD and the change in attenuation from pre-scan to each measured time point is shown in Hounsfield units as mean ± SEM.

### Biodistribution

All mice were sacrificed 24 hours post-injection using carbon dioxide inhalation and were dissected to collect relevant organs. For this study, the heart, lungs, liver, kidneys, spleen, blood, bladder, fecal matter, and tail were isolated, and the remainder of the carcass was collected. Each component was weighed and recorded before being digested in 4 mL of nitric acid for 24 hours at 90°C. 6 mL of deionized water was added to each sample and then they were analyzed using ICP-OES to detect silver content. Data is presented as mean ± SEM.

### Statistical Analysis

All experiments described herein were performed in triplicate, unless otherwise specified. Statistical analysis was performed using GraphPad Prism 9 software with selected statistical tests described in the methods for each experiment.

## Results

### Synthesis and Characterization

Ag_2_S-NP were synthesized using the bulk method (Bulk-Ag_2_S-NP), a commercially available microfluidic mixing chip (CS-Ag_2_S-NP), and SSMS chips with 1 channel (1X-Ag_2_S-NP), 10 parallelized channels (10X-Ag_2_S-NP), and 256 parallelized channels (256X-Ag_2_S-NP). All SSMS chips were monitored during synthesis (Figure S1). While the 1X SSMS chip can produce only 0.1 L/hr of Ag_2_S-NP, the 10X SSMS chip produces Ag_2_S-NP at a rate of 1 L/hr and the 256X SSMS chip produces Ag_2_S-NP at 17 L/hr (Figure 2A,B). In practice, this results in the ability to synthesize 100 mL of Ag_2_S-NP in 1 hour, 6 minutes, or 23 seconds, respectively, demonstrating the possibility of a drastically increased rate of synthesis. The Ag_2_S-NP produced were also found to be consistent in core size and concentration when isolated at various time points throughout the synthesis process (Figure S2).

**Figure 2:**
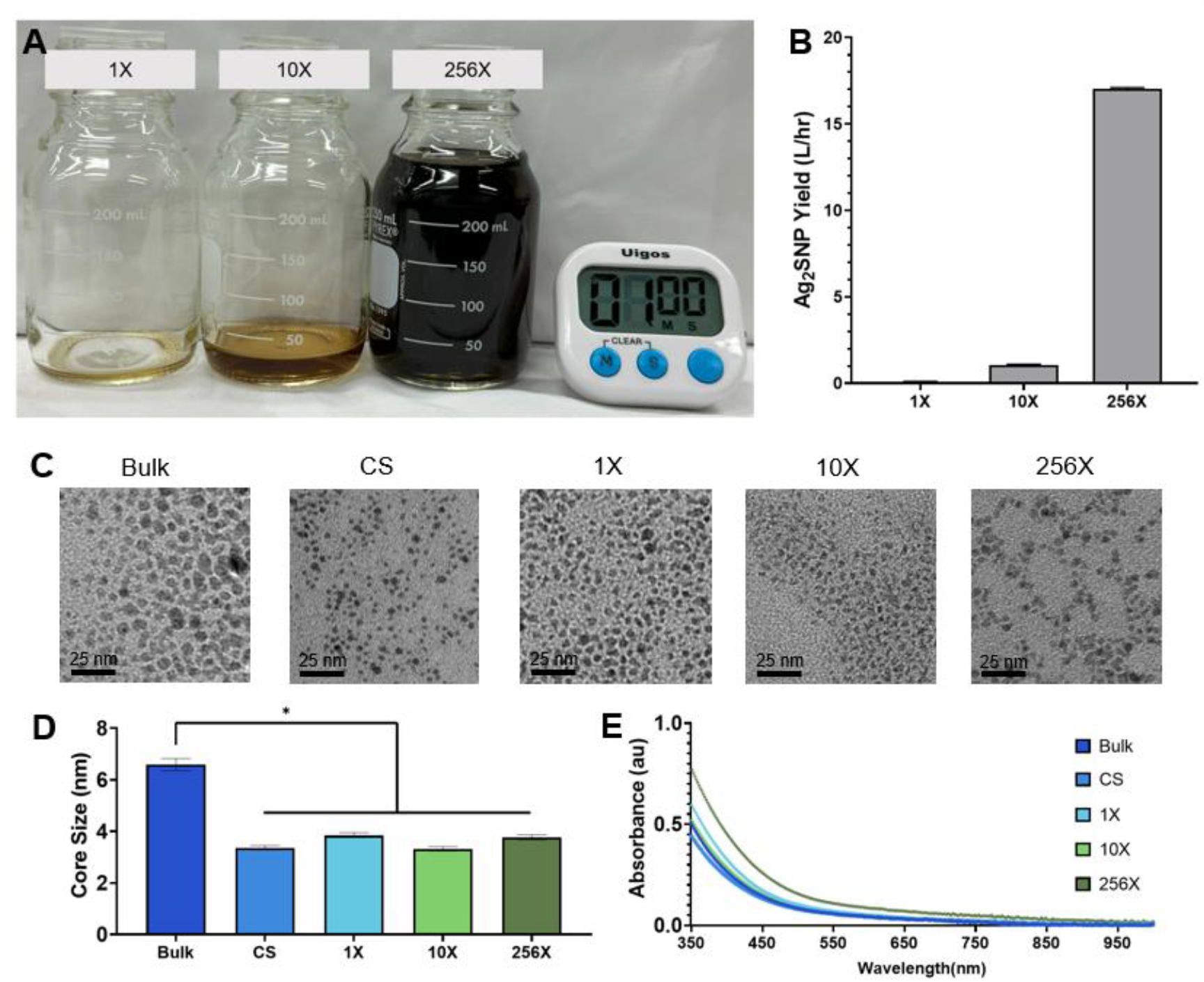
A) Representative images of Ag_2_S-NP produced using each SSMS chip after running for one minute and B) rate of production using each device (mean ± SEM). Characterization of the Ag_2_S-NP synthesized including C) representative TEM micrographs, D) core size measurements (mean ± SEM), and E) UV-visible spectra. Scale bars: 25 nm.

Using TEM, the Bulk-Ag_2_S-NP were found to have a core diameter of 6.6 ± 1.7 nm which is significantly larger than all the Ag_2_S-NP produced with microfluidic chips. The CS-, 1X-, 10X-, and 256X-Ag_2_S-NP were all found to have core diameters of less than 5 nm, indicating that they are all suitable for renal clearance *in vivo*, while the bulk synthetic method does not produce Ag_2_S-NP small enough for this purpose under these conditions. The diameters of the CS-, 1X-, 10X-, and 256X-Ag_2_S-NP were 3.4 ± 0.6 nm, 3.8 ± 0.8 nm, 3.3 ± 0.6 nm, and 3.8 ± 0.7 nm, respectively, and found not to be significantly different (Figure 2C,D). UV-visible absorbance spectra were also very similar for Ag_2_S-NP made via the five synthetic methods, which confirms the formation of Ag_2_S-NP as expected (Figure 2E).^10^ Zeta potential measurements indicated that all of the Ag_2_S-NP samples produced had negative surface charges (Figure S3).

To further scale up the process, we studied the effect of increasing the concentration of the reagents used to further decrease the time needed for synthesis and filtration. This would not only provide a higher production rate, but also increase the reaction product concentration, reducing the total time needed for post-processing. To this end, Ag_2_S-NP were synthesized with standard conditions as well as reagents at 500%, 1000%, and 2000% concentrations. The resulting nanoparticles were all found to be less than 5 nm, in line with the requirement for renal clearance, with no substantial differences in the diameters of the product between the conditions (Figure 3A,B). Additionally, the yield of Ag_2_S-NP was found to scale linearly with increased concentrations, indicating that production does not diminish relative to the increased concentrations (Figure 3C). These results indicate that increased concentrations of reagents may also be a valuable avenue for scaling up the synthesis methods of Ag_2_S-NP.

**Figure 3:**
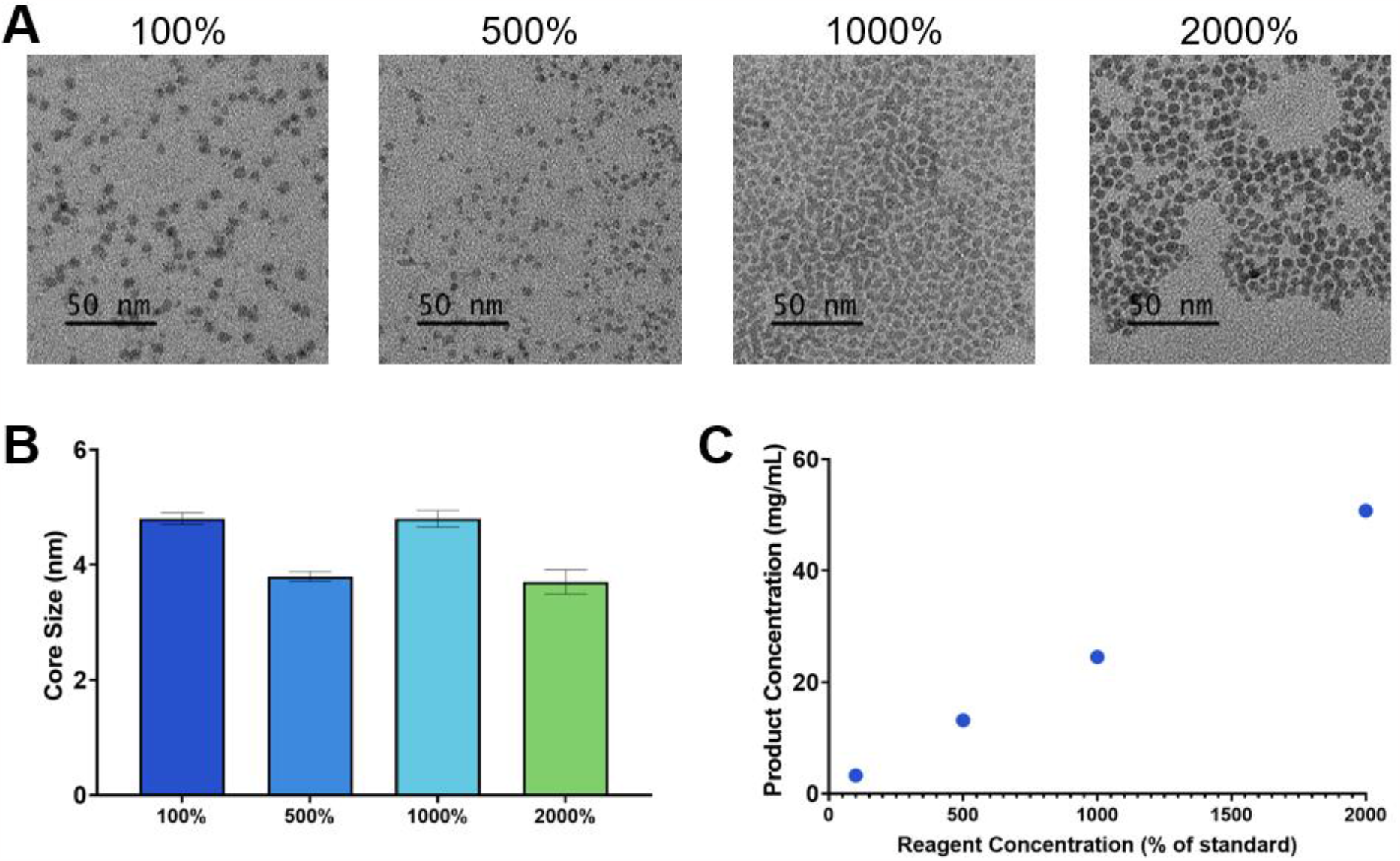
Characterization of the Ag_2_S-NP synthesized with increased concentrations including A) representative TEM micrographs, B) core size measurements (mean ± SEM), and C) product concentration as measured by ICP.

### *In Vitro* Contrast Generation

Computed tomography (CT) scans of the phantoms prepared using each of the Ag_2_S-NP samples synthesized with an SSMS chip revealed no significant difference in attenuation rates as measured by an MI Labs microCT scanner. All three scales of SSMS chips produced Ag_2_S-NP with attenuation rates of about 60 HU·mg/mL. Importantly, the attenuation rates of the SSMS-synthesized Ag_2_S-NP were significantly higher than that of iopamidol, which only had an attenuation rate of 30 HU·mg/mL in this scanner (Figure 4).

**Figure 4:**
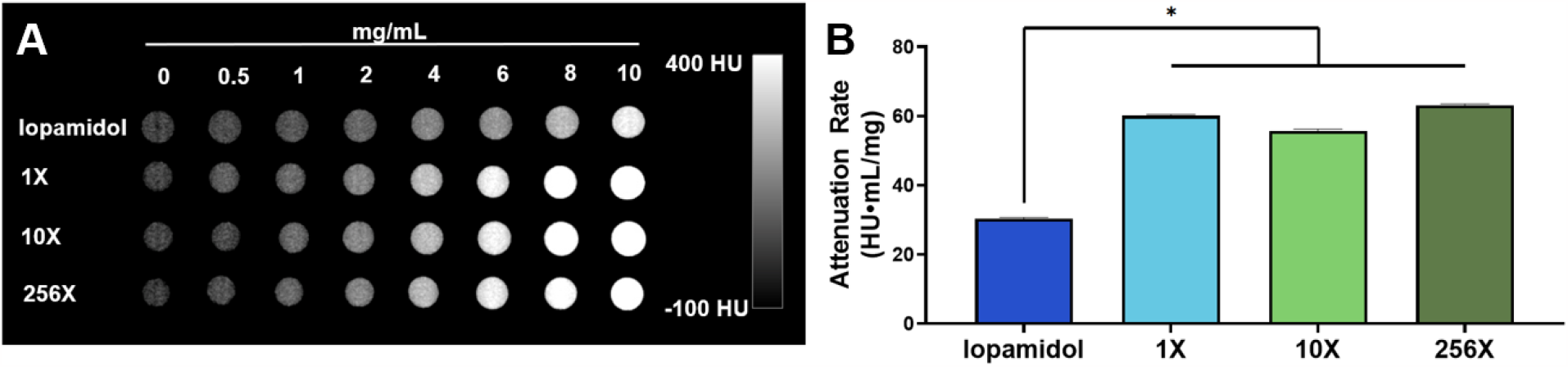
*In vitro* CT imaging. A) Representative μCT scans with iopamidol, 1X-Ag_2_S-NP, 10X-Ag_2_S-NP, 256X-Ag_2_S-NP at concentrations ranging from 0 – 10 mg/mL and B) quantification of the CT attenuation rate for the different solutions (mean ± SD).

### *In Vivo* Imaging

Since the key criterion for clinical translation of an imaging agent containing a heavy element such as silver is excretion, we studied their renal clearance in mice via CT imaging. After injection with Ag_2_S-NP, CT scans revealed that Ag_2_S-NP prepared with each type of microfluidic chip was similarly excreted primarily through the renal clearance route. Visibly identifiable contrast is present in the kidneys and bladder beginning only 5 minutes after injection, continues to be noticeable through the 2-hour time point, but is mostly eliminated by the 24-hour scan (Figure 5A, S4). Quantitative analysis of the scans confirms that increased contrast is present in the kidneys and bladder at 2 hours post injection, but mostly eliminated by 24 hours (Figure 5B,C). Virtually no contrast was observed in the heart or liver (Figure S5). These results were consistent across the CS-Ag_2_S-NP, 1X-Ag_2_S-NP, 10XAg_2_S-NP, and 256X-Ag_2_S-NP, and the variation in results can be attributed to varying times of excretion for each mouse.

**Figure 5:**
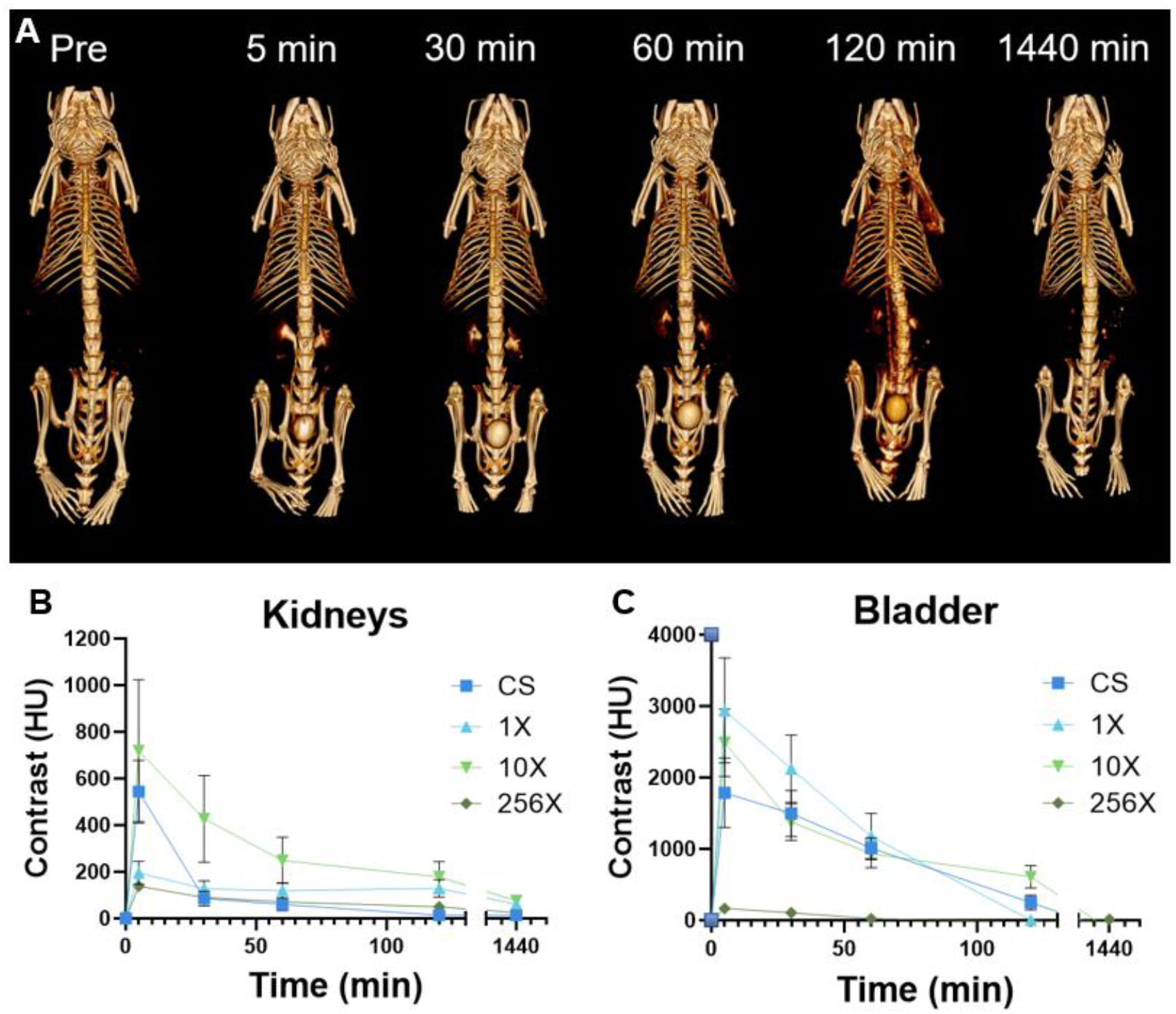
A) Representative 3D μCT images showing Ag_2_S-NP being renally cleared. Images include pre-injection and 5 min, 30 min, 60 min, 120 min, and 1440 min post-injection. Quantification of CT attenuation in the B) kidneys and C) bladder at each time point. n=5 per group. Data is presented as mean ± SEM.

### Biodistribution

24 hours after the intravenous injection of Ag_2_S-NP, mice were sacrificed and their organs were collected for analysis with ICP-OES. Quantitative analysis of the silver content in each collected organ revealed that there was very low retention of these nanoparticles, indicating that they were extensively renally cleared. The highest %ID was found in the spleen and kidneys, indicating primary renal clearance, with very small amounts found in RES organs. Carcass, including the tail, retention can be primarily attributed to the injection method (Figure S6). Importantly, all synthetic methods resulted in around 90% clearance of the injected dose within 24 hours (Figure 6).

**Figure 6:**
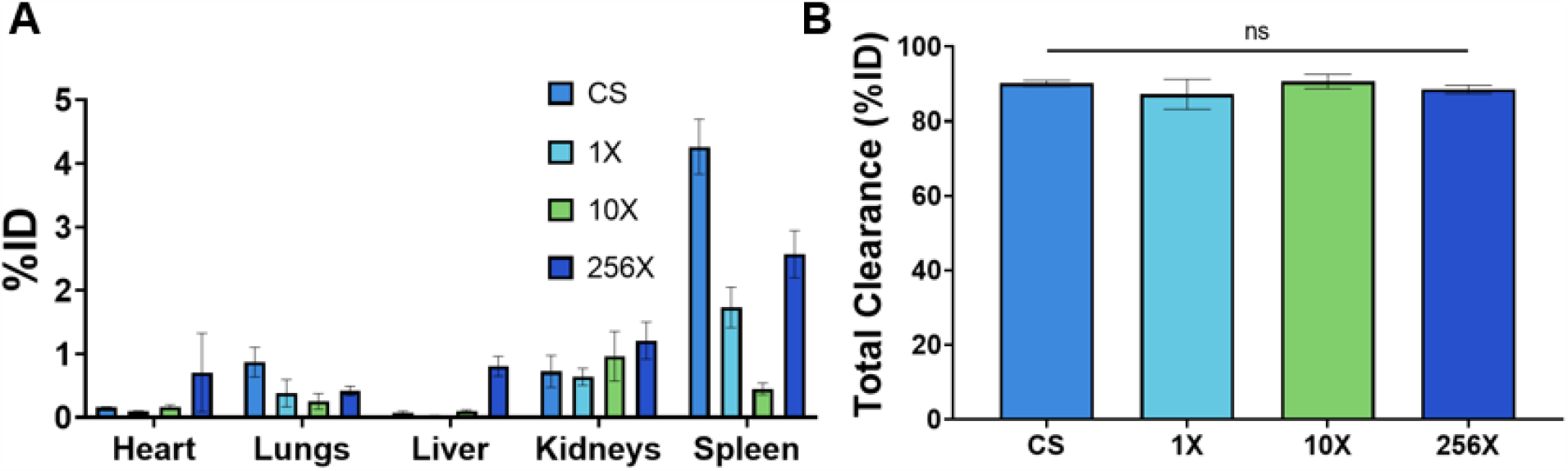
A) Biodistribution of Ag_2_S-NP in mice 24 hours post-injection. n=5 per group. Data is presented as mean ± SEM. B) Total clearance of Ag_2_S-NP in mice within 24 hours. Data is presented as mean ± SEM.

## Discussion

The results presented herein demonstrate a drastic capacity for increased synthesis without sacrificing particle quality, tunability, or imaging contrast generation. Compared to the more than 387 hours needed to produce one human dose of the Ag_2_S-NP with a single channel microfluidic chip, using the 256X SSMS, one dose could be produced in 2.3 hours. Coupled with 2000% concentration of reagents used, which was also demonstrated to be an effective method for increasing production, the synthesis could be completed in less than 7 minutes, representing scale-up of 3400-fold. These combined scale-up efforts could result in the production of > 5 doses per hour by a single chip. This impressive increase creates a feasible path for clinical level synthesis of Ag_2_S-NP, since additional avenues for further production increases remain, such as increasing the parallelization of the chip (e.g. to 1000 channels), running multiple chips at once and increasing flow rates.

Our synthesis methods using SSMS chips consistently resulted in sub-5 nm nanoparticles. Moreover, size can be further tuned by adjusting the relative flow rates of the reagents.^10^ The contrast generation rates of the resulting Ag_2_S-NP are substantially improved over current clinical agents. We demonstrated potent *in vivo* contrast generation compared to commercial standards and a high safety profile via renal clearance. This points to the clinical potential of Ag_2_S-NP and the possibility of its use in place of iodinated agents in some applications.^36^ For example, implementation of Ag_2_S-NP in the clinical paradigm of breast cancer imaging can provide better screening and earlier detection in high-risk populations, such as those with dense breast tissue.^37, 38^ Furthermore, this low-cost agent will be easy to implement due to its compatibility with imaging modalities that are readily available in the clinic such as mammography.

This study has also confirmed that the sub-5 nm Ag_2_S-NP produced using microfluidic synthesis methods have best-in-class renal clearance.^8^ With a 24-hour clearance of about 90% of injected dose for all microfluidic systems tested, Ag_2_S-NP have some of the highest clearance rates compared to other similarly sized metallic nanoparticles.^39^ The clearance results displayed here are consistent with previous work with Ag_2_S-NP evaluating their clearance, which found an 85% to 93% clearance rate in 24 hours.^8, 10^

Across each metric evaluated here, i.e. nanoparticle size, absorption spectra, imaging contrast *in vitro* and *in vivo*, and biodistribution, each SSMS synthesis platform has been proven to result in nanoparticles of consistent quality and performance. This confirms the potential for this system to be used for scaled up production of Ag_2_S-NP and other inorganic nanoparticles. Further, since the system described here was developed in accordance with industry standards for pharmaceutical synthesis, the transition to mass production will be easier. This study removes barriers to large animal studies and the eventual clinical use of Ag_2_S-NP in an innovative and potentially widely applicable manner.

One limitation to this study is the microfluidic chips used were not run to exhaustion, and therefore claims cannot be made about the longevity or reusability of the chips past the syntheses specifically described. However, when similar chips were used for lipid nanoparticle production, it was shown that they could be reused multiple times without compromising the product.^29^ Additionally, though this work tests up to 256 parallelized channels and reagent concentrations increased up to 2000% of the original, higher throughput methods could still be explored, such as higher flow rates, although this can impact the characteristics of the resulting nanoparticles.^40^ Furthermore, although it is expected that similar results would be observed with other inorganic nanoparticles, this has yet to be explored with the SSMS system. Finally, the contrast generation and biodistribution profile presented are limited to the μCT scanner and mouse model used.

In the future, it may be advantageous to develop a system for synthesis that includes formulation, filtration, and concentration of nanoparticles in one continuous system. To this end, it would be beneficial to evaluate the feasibility of incorporating a tangential flow filtration system in place of the current system of centrifugation with centrifugal filters. Further studies of the Ag_2_S-NP biodistribution, clearance, and safety could also be performed in large animal models, as the high-capacity synthesis methods presented herein have made these studies more feasible.

## Conclusion

The SSMS system can provide large-scale increase in production of Ag_2_S-NP without sacrificing nanoparticle quality or imaging utility. We demonstrated total production rates up to 17 L/hr when using a 256X channel chip, which is more than 100-fold higher than a single channel chip. This production increase could be further increased by use of multiple chips or 1000X channel chips, for example. Moreover, when combined with increased concentration of reagents (e.g. as much as 20x), we were able to synthesize Ag_2_S-NP on a 3,400-fold scale. Additionally, Ag_2_S-NP produced using this novel system retain their contrast and clearance properties *in vivo*. The best-in-class clearance rates of the Ag_2_S-NP, paired with significantly improved contrast generation compared to clinically available iodinated agents, make this contrast agent promising for further development and confirms the need for clinical scale production capacity. This study validates the potential for a substantial scale up in production, resulting in an improved outlook for clinical translation for Ag_2_S-NP.

## Acknowledgements

The studies presented here were completed with funding from the NIH via R01 CA227142 (DPC), F31-EB034165 (DNRB) and R25 CA140116, which supported NO. This work was also partially supported by the NSF GRFP (SJS) and the Brody Family Medical Trust Fund Fellowship (JCH). This work was partially completed in the Singh Center for Nanotechnology as part of the NSF National Nanotechnology Coordinated Infrastructure Program under grant NNCI-2025608.

The authors would also like to acknowledge the Electron Microscopy Resource Lab at the University of Pennsylvania, and specifically Sudheer Molugu, for assistance with TEM. They would also like to thank Eric Blankemeyer for his assistance with μCT scanning and Dr. David Vann and Dr. David Burney for their assistance with ICP.

## Notes

### Competing Interest Statement

The authors have declared no competing interest.

