## Supplemental Data for "Towards the Clinical Translation of a Silver Sulfide Nanoparticle Contrast Agent: Large Scale Production with a Highly Parallelized Microfluidic Chip"

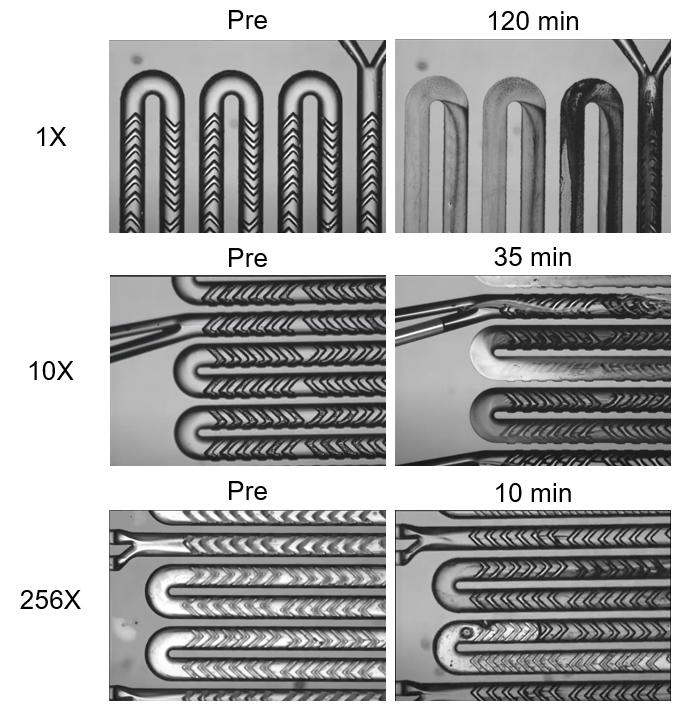

Figure S1. Images of SSMS chips in use over time.

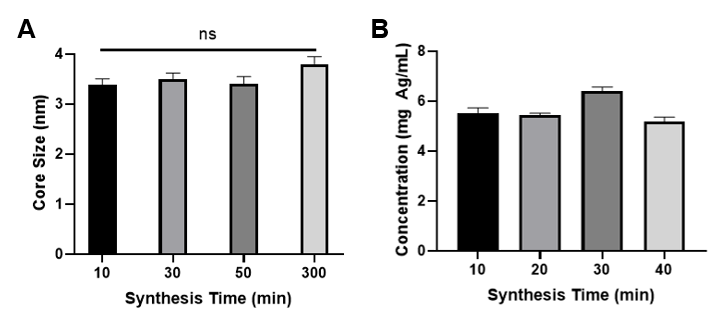

Figure S2. Ag_2_S-NP A) product size as evaluated by TEM and B) concentration as evaluated by ICP measured over time during synthesis by a 1X SSMS chip.

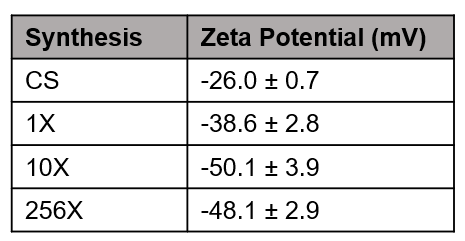

Figure S3. Table of zeta potential values for each Ag_2_S-NP synthetic condition.

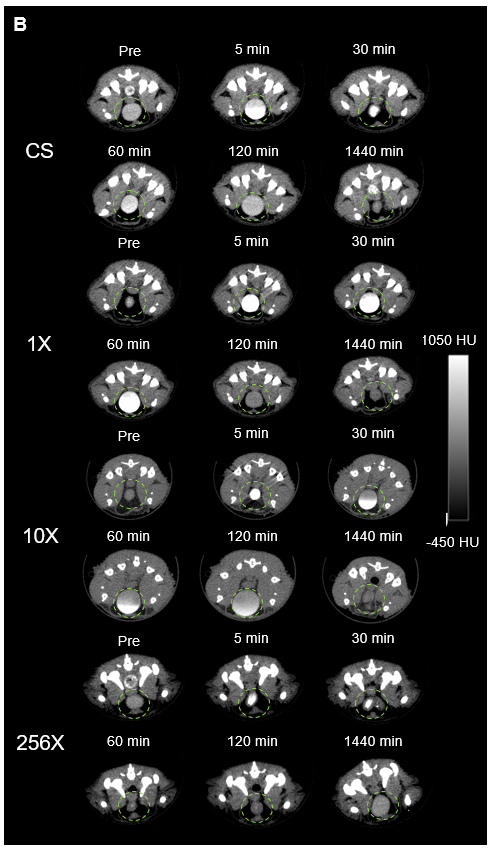

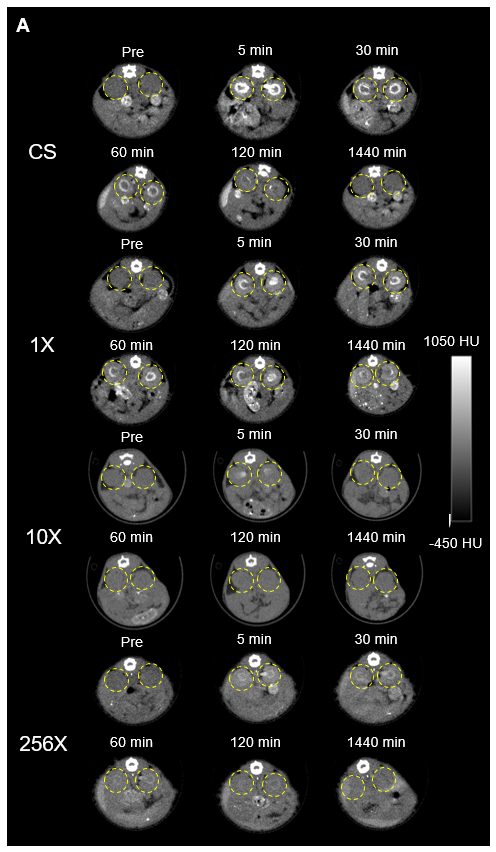

Figure S4. Representative µCT images showing Ag_2_S-NP being renally cleared through A) the kidneys (yellow circle) and B) the bladder (green circle).

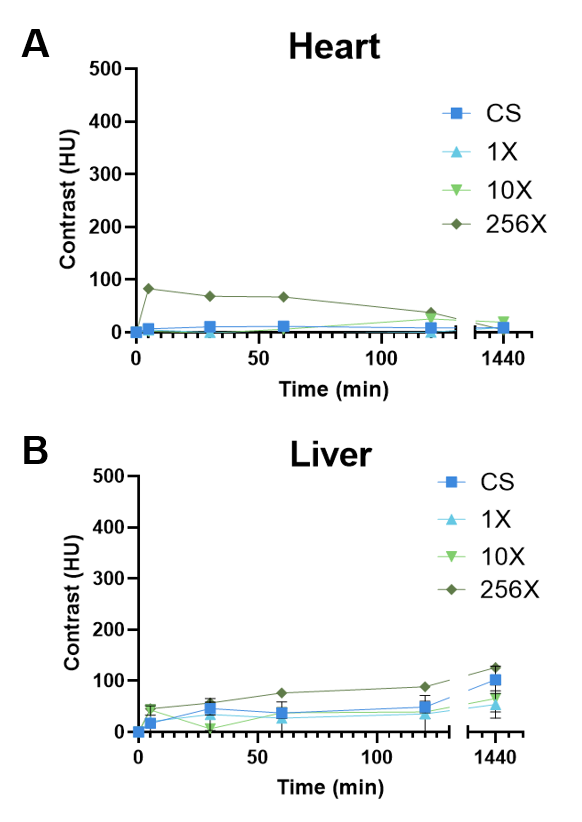

Figure S5. Quantification of CT attenuation in the A) heart and B) liver at each time point. n=5 per group. Data is presented as mean ± SEM.

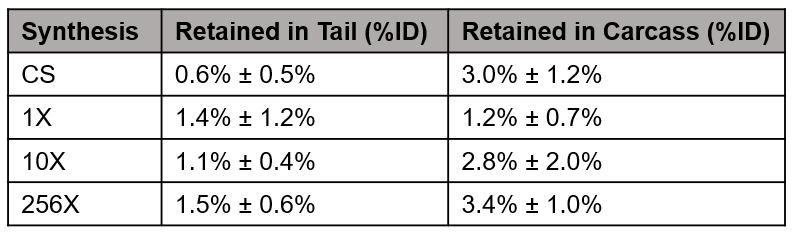

Figure S6. Silver content retained by the carcass and the tail of mice injected with Ag_2_SNP.
